# Mechanistically Informed Machine Learning Links Non-Canonical TCA Cycle Activity to Warburg Metabolism and Hallmarks of Malignancy

**DOI:** 10.1101/2025.08.01.668082

**Authors:** Lin Lihao, Lapi Francesco, Bruno G. Galuzzi, Marco Vanoni, Lilia Alberghina, Chiara Damiani

## Abstract

Cancer cells undergo extensive metabolic rewiring to support growth, survival, and phenotypic plasticity. A non-canonical variant of the tricarboxylic acid (TCA) cycle, characterized by mitochondrial-to-cytosolic citrate export, has emerged as critical for embryonic stem cell differentiation. However, its role in cancer remains poorly understood.

Here, we present a two-step computational framework to systematically analyze the activity of this non-canonical TCA cycle across over 500 cancer cell lines and investigate its role in shaping hallmarks of malignancy. First, we applied constraint-based modeling to infer cycle activity, defining two complementary metrics: *Cycle Propensity*, measuring the likelihood of its engagement in each cell line, and *Cycle Flux Intensity*, quantifying average flux through the reaction identified as rate-limiting. We identified distinct tumor-specific patterns of pathway utilization. Notably, cells with high Cycle Propensity preferentially rerouted cytosolic citrate via aconitase 1 (ACO1) and isocitrate dehydrogenase 1 (IDH1), promoting *α*-ketoglutarate (*α*KG) and NADPH production. Elevated engagement of this cycle strongly correlated with Warburg-like metabolic shifts, including decreased oxygen consumption and increased lactate secretion.

In the second step, to uncover non-metabolic transcriptional signatures associated with non-canonical TCA cycle activity, we performed machine learning–based feature selection using ElasticNet and Random Forest, identifying robust gene signatures predictive of cycle activity. Over-representation analysis revealed enrichment in genes involved in invasiveness, angiogenesis, stemness, and key oncogenic pathways. Analysis of DepMap gene dependency data revealed that TCA cycle activity correlates with differential vulnerability to perturbation of these oncogenic pathways, reinforcing the functional relevance of identified transcriptional signatures. To further interpret the predictive models, SHapley Additive exPlanations (SHAP) was applied to prioritize genes contributing most to non-canonical cycle activity, suggesting novel candidates for experimental investigation.

Overall, our framework enables comprehensive analysis of non-canonical TCA cycle dynamics and uncovers potential links between metabolic plasticity and malignant phenotypes.

## 1 Introduction

A non-canonical variant of the TCA cycle was recently described by Arnold et al. [1] as a cross-compartment metabolic pathway that links mitochondrial and cytosolic metabolism, hereinafter referred to as the Arnold Cycle. Unlike the classical TCA cycle confined to mitochondria, the Arnold Cycle involves the export of citrate to the cytosol via the SLC25A1 transporter, its cleavage by ATP citrate lyase (ACLY) into acetyl-CoA and oxaloacetate, and the subsequent reduction of oxaloacetate to malate by cytosolic malate dehydrogenase (MDH1) (*Fig. 1a*).

**Figure 1:**
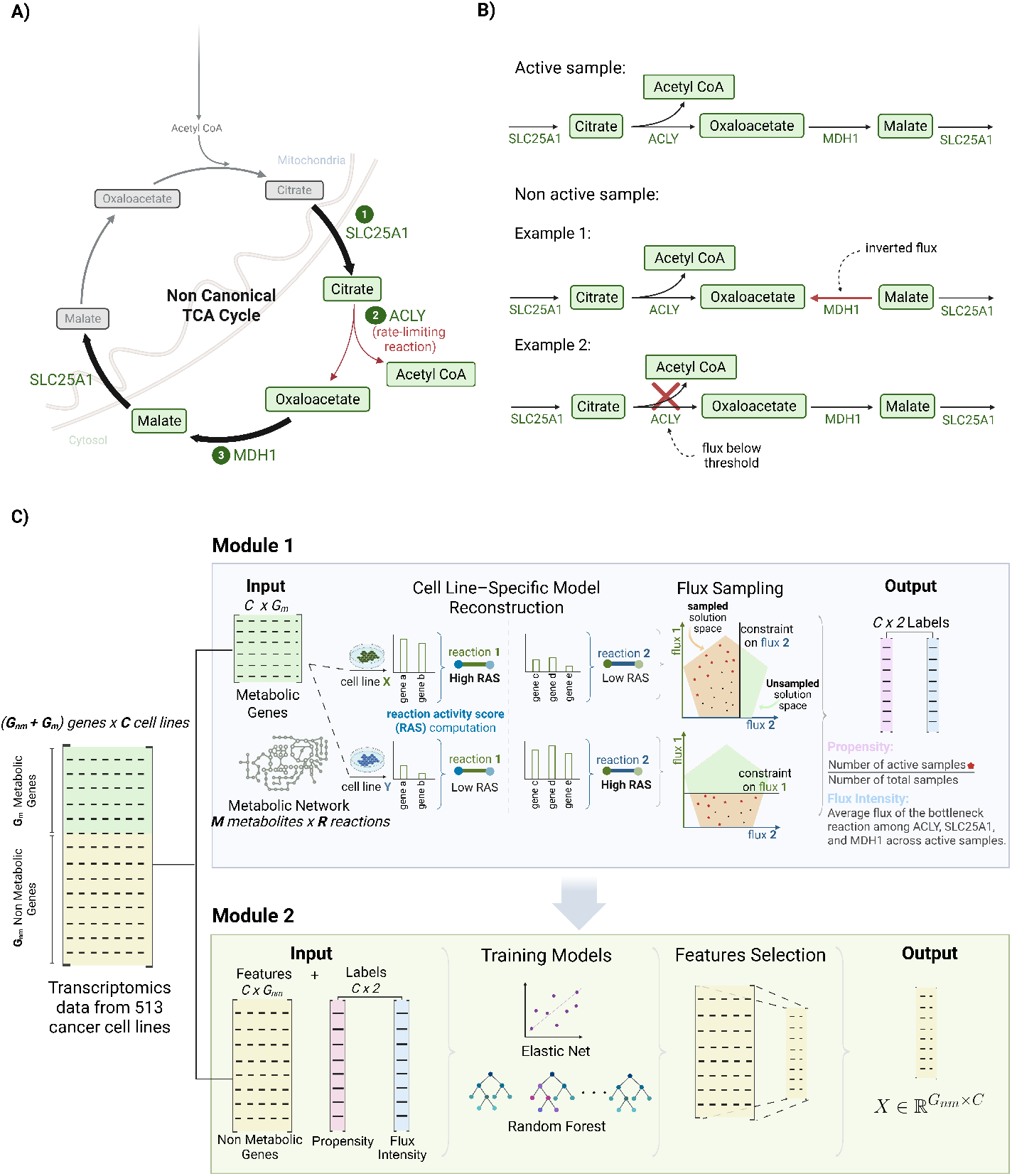
Schematic representation of the non-canonical TCA cycle and the computational pipeline. **A)**, The non-canonical TCA cycle involves: (1) export of mitochondrial citrate to the cytosol via SLC25A1; (2) cleavage of citrate by atp citrate lyase (ACLY) to produce acetyl-CoA and oxaloacetate; (3) reduction of oxaloacetate to malate by cytosolic malate dehydrogenase (MDH1), which can be re-imported into the mitochondria via SLC25A1, thus completing the cycle. Arrow thickness reflects the average flux value of each reaction across active samples. ACLY is identified as the bottleneck reaction. **B)**,Definition of active and inactive samples based on flux thresholds and directionality. Samples are classified as *active* when all three hallmark reactions of the cycle—`SLC25A1` →`ACLY` → MDH1—carry flux above a minimal threshold (10^−6^) and operate in the expected direction. In contrast, *inactive* samples include cases where at least one reaction operates in the opposite direction (Example 1), or where one reaction, such as `ACLY`, carries sub-threshold flux, failing to sustain the cycle (Example 2). **C)** Computational pipeline: Module 1 integrates metabolic gene expression (*G*_*m*_) with a constraint-based model to estimate pathway activity (*Cycle Propensity* and *Flux Intensity*). These estimates serve as labels for Module 2, where machine learning models are trained on non-metabolic gene expression (*G*_*nm*_) to select transcriptional predictors of pathway activity.

Engagement of this cycle bas been reported in both normal and cancer cells and has been linked to transitions in cellular identity [1]. In embryonic stem cells, for instance, it is required to exit from naïve pluripotency, and its inhibition prevents proper differentiation.

While its importance in stem cell biology is increasingly recognized, the regulation and function of Arnold Cycle in cancer remain poorly understood. Whether it merely reflects a metabolic adaptation or contributes more broadly to malignant traits such as transcriptional plasticity or invasiveness is still unclear.

To shed light on how cancer cells exploit metabolic circuits to support phenotypic plasticity and progression, we aim to systematically characterizate the transcriptional programs associated with Arnold Cycle activity across diverse tumor types. However, direct measurements of compartment-specific metabolic fluxes are lacking in most cancer datasets, making it difficult to infer the activity of pathways that span multiple cellular compartments and require directional consistency, as in the Arnold Cycle. In this context, constraint-based modeling offers a particularly well-suited computational approach to estimate intracellular fluxes from gene expression profiles [2]. Nevertheless, these methods are inherently limited to metabolic genes and fail to capture regulatory or downstream transcriptional programs outside the metabolic network [3, 4]. To overcome the above limitations, we developed a mechanistically informed machine learning (ML) framework that integrates constraint-based metabolic modeling with supervised learning. As schematized in *Fig. 1c*, we first applied constraint-based modeling to derived quantitative metrics of Arnold Cycle activity across cancer cell lines. We then used these metrics as labels in a supervised learning problem to select transcriptional features predictive of Arnold Cycle activity. We applied our framework to over 500 cancer cell lines, revealing robust transcriptional programs associated with Arnold Cycle activity. Beyond serving as labels for machine learning, our *in silico* metrics also uncovered previously unrecognized connections between Arnold Cycle activity and broader patterns of metabolic reprogramming in cancer, offering insights into how cancer cells exploit metabolic flexibility to sustain aggressive phenotypes.

## 2 Results

### 2.1 Tumor-Specific Usage and Metabolic Rerouting of the Arnold Cycle

In the first module of our two-step computational strategy (*Fig. 1c*), we inferred the activity of the Arnold Cycle across 513 cancer cell lines. To this aim, we used ENGRO2.2, an updated version of the core metabolic network ENGRO2 [2].

For each line, we reconstructed a cell-specific metabolic model constrained by gene expression data and sampled feasible steady-state flux distributions. From these samples, we derived two complementary metrics: *Cycle Propensity*, defined as the fraction of sampled states in which the three core reactions (SLC25A1, ACLY, MDH1) of the Arnold Cycle are simultaneously active; and *Cycle Flux Intensity*, measuring the average flux through the cycle’s bottleneck reaction in active configurations (*Fig. 1b*).

Arnold Cycle activity displays marked heterogeneity across tumor types, revealing distinct lineage-specific patterns (*Fig. 2a*). Lung, stomach, breast, and pancreatic tumors displayed wide variability in both *Cycle Propensity* and *Cycle Flux Intensity*, indicating substantial intra-lineage diversity. In contrast, skin, lymphoid, and esophageal tumors formed relatively compact clusters characterized by high *Cycle Propensity* but low *Flux Intensity*. By comparison, central nervous system and myeloid tumors exhibited intermediate levels of both *Cycle Propensity* and *Flux Intensity*, suggesting moderate activity of the Arnold Cycle in these lineages.

**Figure 2:**
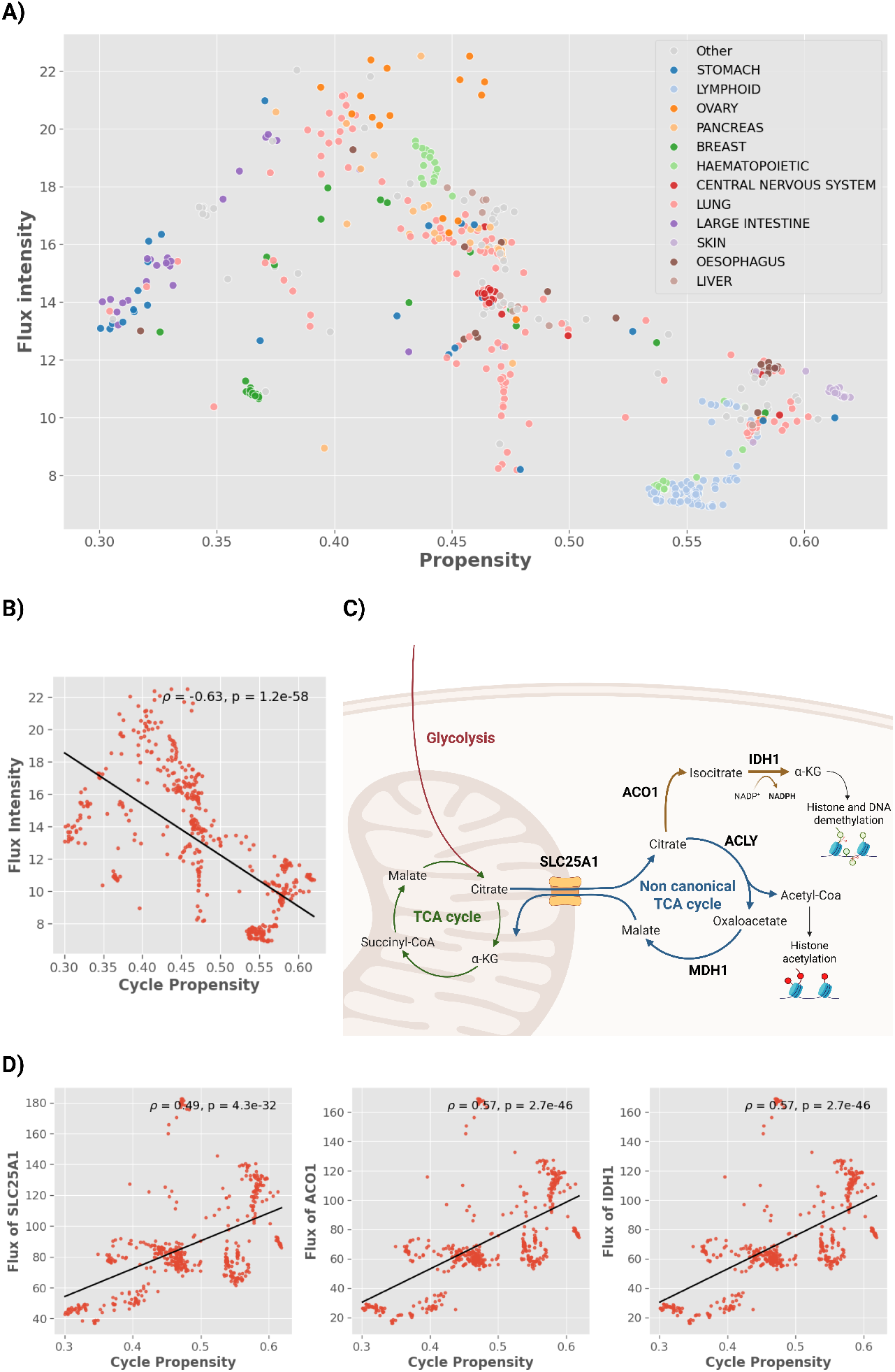
Heterogeneous usage and routing of the non-canonical TCA cycle in cancer cell lines. **(A)** Scatterplot of *Cycle Propensity* versus *Flux Intensity* across 513 cancer cell lines, color-coded by tissue type, reveals lineage-specific patterns and intra-lineage heterogeneity. Tumor types with fewer than 10 cell lines were grouped and displayed in grey. **(B)** *Cycle Propensity* and *Flux Intensity* are negatively correlated (*ρ* = − 0.63, *p* = 1.25 ×10^−58^). **(C)** Schematic of cytosolic citrate routing, illustrating the bifurcation between ACLY-mediated acetyl-CoA production and the ACO1–IDH1 axis generating *α*KG and NADPH. **(D)** Positive correlations between *Cycle Propensity* and flux through SLC25A1 (*left*), ACO1 (*mid*), and IDH1 (*right*) support preferential oxidative routing in high-propensity cell lines.

Although *Cycle Propensity* and *Cycle Flux Intensity* were defined to capture complementary aspects of Arnold Cycle behavior, they were found to be negatively correlated (*ρ* = −0.63, *p* = 1.25 *×* 10^−58^; *Fig. 2b*). This inverse relationship suggests that a higher likelihood of Arnold Cycle engagement does not necessarily correspond to increased metabolic throughput. To gain mechanistic insight into this unexpected anticorrelation, we considered that *Cycle Flux Intensity* was defined based on the flux through the pathway’s rate-limiting step. We therefore examined which reaction constrained flux across all cell lines. In all models, ACLY was consistently identified as the bottleneck, catalyzing the conversion of cytosolic citrate into acetyl-CoA and oxaloacetate.

Given this observation, we next investigated how citrate flux is redistributed in cell lines with higher *Cycle Propensity*. As an initial step, we asked whether increased *Cycle Propensity* coincides with enhanced mitochondrial export of citrate. Indeed, we found a significant positive correlation between *Cycle Propensity* and flux through SLC25A1—the mitochondrial citrate transporter (*ρ* = 0.49, *p* = 4.26 × 10^−32^; *Fig. 2d*)—suggesting that greater pathway engagement is associated with increased cytosolic citrate export. However, rather than being channeled through ACLY—the rate-limiting step—this exported citrate appears to be rerouted toward alternative cytosolic pathways. To identify these pathways, we focused on the alternative branch that use cytosolic citrate as a substrate in the ENRO2.2 network, namely IDH. Indeed, the flux through IDH1 - which converts isocitrate to *α*KG - showed a strong positive correlation with Cycle Propensity (*ρ* = 0.57, *p* = 2.69 × 10^−46^; *Fig. 2d*). This flux is preceded by the activity of ACO1, which converts citrate to isocitrate, suggesting a rerouting of cytosolic citrate toward oxidative metabolism via the ACO1–IDH1 axis (*Fig. 2c*). To specifically capture this rerouting in the context of Arnold Cycle engagement, flux values for SLC25A1, ACO1, and IDH1 were computed exclusively from sampled states in which the cycle was active.

Together, these findings suggest that in cell lines exhibiting a preference for Arnold Cycle engagement, cytosolic citrate may not be exclusively funneled through ACLY for acetyl-CoA production. Instead, a substantial portion may be diverted toward oxidative reactions mediated by IDH1, a well-known source of cytosolic *α*KG and NADPH [5, 6]. This rerouting, combined with the persistent bottleneck at ACLY, provides a mechanistic explanation for the observed anticorrelation between *Cycle Propensity* and *Cycle Flux Intensity* and highlights how the Arnold Cycle represents only one configuration within a more flexible program of metabolic shift that also includes oxidative branching via ACO1 and IDH1.

These findings also underscore the importance of jointly interpreting both metrics. While *Cycle Flux Intensity* captures the effective flux through the ACLY-limited Arnold Cycle, *Cycle Propensity* reflects a broader transcriptional predisposition toward cytosolic citrate metabolism, including alternative routing through ACO1 and IDH1. Considering both metrics together thus yields a more comprehensive view of this metabolic rewiring.

### 2.2 Non-canonical TCA Cycle Activity Associates with Warburg-like Metabolism

Proliferating cells frequently adopt the Warburg effect—a metabolic reprogramming where glycolysis dominates over oxidative phosphorylation even under oxygen-rich conditions, resulting in elevated lactate secretion [7, 8]. This adaptation fuels rapid biomass generation by providing energy and biosynthetic precursors [9]. To investigate whether non-canonical TCA cycle engagement influences this metabolic state, we analyzed its relationship with two Warburg hallmarks: oxygen consumption and lactate production.

For each cell line, we computed average oxygen uptake and cytosolic lactate export fluxes across all sampled steady-state flux distributions. Strikingly, higher non-canonical *Cycle Propensity* strongly correlated with reduced oxygen utilization (*ρ* = 0.79, *p* = 5.50 < 10^−91^) and increased lactate secretion (*ρ* = 0.74, *p* < *p* = 6.24 < 10^−110^), despite unconstrained oxygen availability in simulations (*Fig. 3a*).

**Figure 3:**
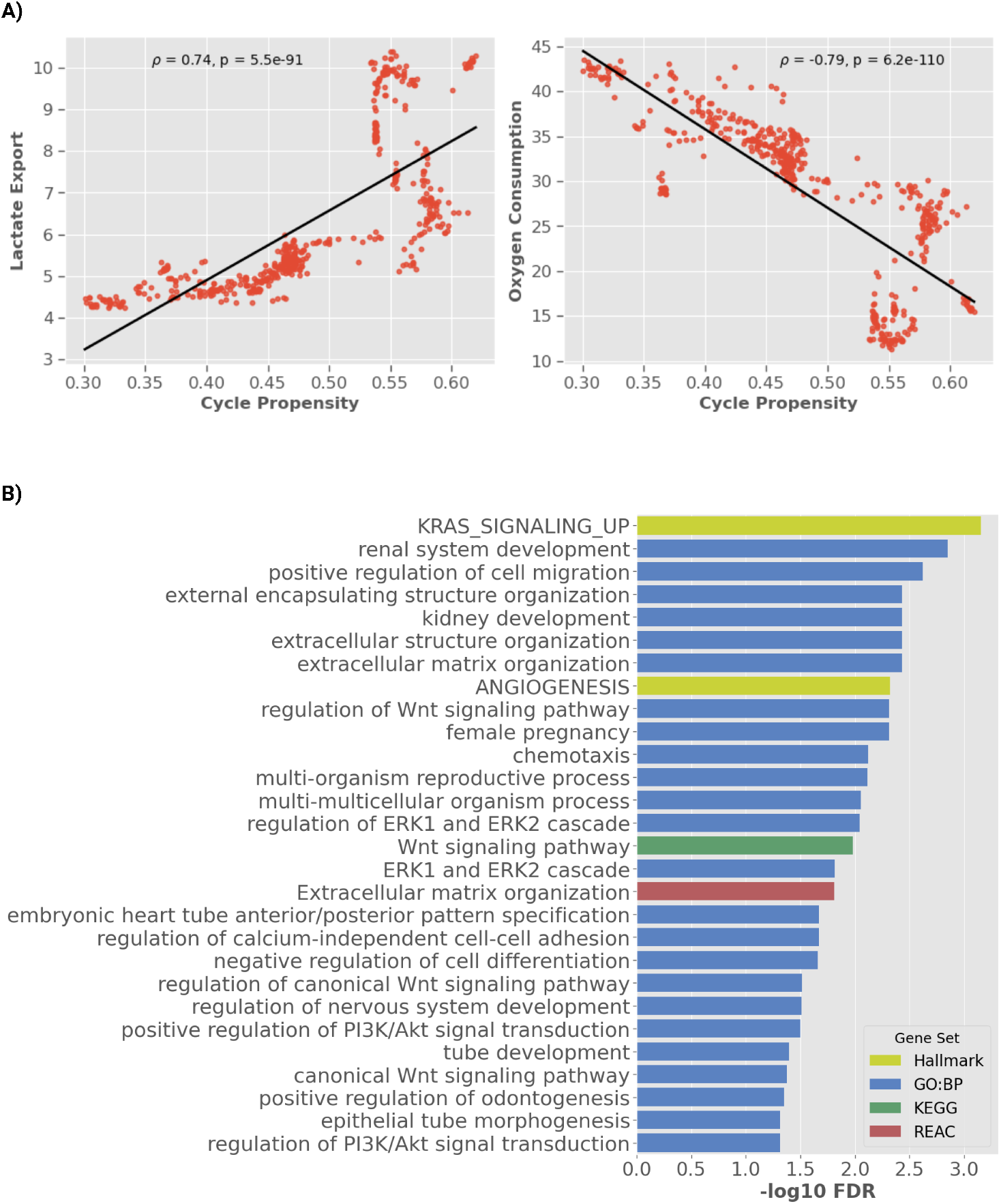
Non-canonical TCA cycle activity links with Warburg metabolism and pro-malignant transcriptional programs. **(A)** Relationship between *Cycle Propensity* and key Warburg metabolism markers across cancer cell lines, evaluated using Pearson correlation. (*left*) *Cycle Propensity* positively correlates with lactate secretion (*ρ* = −0.74, *p* = 5.5 × 10^−91^). (*right)* A strong negative correlation is observed between Cycle Propensity and oxygen consumption (*ρ* = 0.79, *p* = 6.2 × 10^−110^), despite unconstrained oxygen availability in the simulations. **(B)** Over-representation analysis of 314 genes predictive of *Cycle Propensity* and *Flux Intensity*, selected through ElasticNet and Random Forest regressors. Enriched gene sets are derived from GO Biological Process (GO:BP), KEGG, and Reactome (REAC) databases. Bars represent enrichment significance expressed as − log_10_(FDR), color-coded by database.

These findings are consistent with the hypothesis that non-canonical TCA cycle activity is associated with or contributes to a Warburg-like metabolic phenotype. In line with this, recent models propose that its activation promotes biosynthetic precursor flux while reducing oxidative metabolism, reinforcing this metabolic state [10].

### 2.3 Non-Canonical TCA Cycle Activity Defines a Malignancy-Linked Transcriptional Landscape

To uncover non-metabolic transcriptional signatures associated with non-canonical TCA cycle activity, in the second module of our two-step computational strategy *(Fig. 1c)*, we trained regression models to predict both *Cycle Propensity* and *Cycle Flux Intensity* from gene expression profiles, using a multi-target approach. The resulting models achieved high predictive performance across cross-validation, with *R*^2^ about 0.9 for both outputs (*Supplementary Tab. S3 S4*). Feature selection leveraged ElasticNet and Random Forest regressors to capture complementary linear and non-linear dependencies. For each method, we prioritized genes consistently selected across multiple cross-validation folds (Methods 4.7.3) and combined these to derive a comprehensive panel of 314 genes (*Supplementary Tab. S1*).

We first evaluated the functional coherence among these genes through protein–protein interaction (PPI) network analysis. Of the 314 genes, 268 are mapped to the PPI network, forming 80 edges—significantly more than the 52 edges expected by random chance (*p* = 2.21 × 10^−4^); the full network map is provided in *Supplementary Fig S1*. The resulting network showed moderate connectivity, with an average node degree of 0.597. Subsequent clustering analysis highlighted one prominent cluster containing 41 genes, accompanied by several smaller clusters. Notably, the major cluster identified *HRAS*—a central component of the RAS signaling cascade—as the primary hub *(Supplementary Fig. S2)*, underscoring its potential role as a key integrator of the transcriptional interactions associated with non-canonical TCA cycle activity.

To systematically explore the biological roles of the 314 selected transcriptional features, we performed Over-Representation Analysis (ORA; *Fig. 3b*). The selected genes were tested for enrichment across GO Biological Processes, KEGG, Reactome, and MSigDB Hallmark gene sets. The background set included 32,555 genes, excluding pseudogenes and canonical metabolic genes. Statistical significance was assessed using a hypergeometric test with Benjamini–Hochberg correction (*FDR*< 0.05).

The selected genes showed strong enrichment for processes involved in tumor progression. Among the most significant were *positive regulation of cell migration* (*FDR* = 2.378 × 10^−3^) and *extracellular matrix organization* (*FDR* = 3.698 × 10^−3^). These findings suggest a shift in gene expression toward a more invasive tumor phenotype.

We also observed enrichment for angiogenesis-related processes (*FDR* = 4.775 × 10^−3^). To the best of our knowledge, no direct link has yet been experimentally established between enzymes involved in the non-canonical TCA cycle and angiogenesis in cancer, highlighting a potentially unexplored connection.

Several key oncogenic signaling pathways were also enriched. These included *regulation of Wnt signaling* (*FDR* = 4.874 × 10^−3^), *PI3K/AKT signaling* (*FDR* = 3.146 × 10^−2^), *ERK1/2 cascade* (*FDR* = 9.100 × 10^−3^), a core component of the MAPK pathway, and *KRAS signaling up* was significantly overrepresented (*FDR* = 6.963 10^−4^). These pathways are recurrently implicated in multiple hallmarks of cancer [11, 12, 13, 14], and their enrichment suggests that non-canonical TCA cycle activity both influences and is shaped by diverse oncogenic signaling cascades. Given the pan-cancer nature of our analysis, this observation points to a heterogeneous and context-dependent metabolic–signaling interplay that may contribute to distinct tumor phenotypes across lineages.

Finally, we observed enrichment in processes related to development and differentiation, such as *kidney system development* (*FDR* = 3.698 × 10^−3^), *regulation of nervous system development* (*FDR* = 3.076 × 10^−2^), and *negative regulation of cell differentiation* (*FDR* = 2.177 × 10^−2^). These transcriptional patterns are indicative of a stem-like phenotype and are consistent with previous evidence implicating non-canonical TCA cycle activity in the regulation of stem cell fate [1].

Collectively, these results reveal a robust transcriptional signature associated with non-canonical TCA cycle activity, including key hallmarks of malignancy, including programs linked to invasion, stemness, angiogenesis, and oncogenic signaling.

### 2.4 SHAP Highlights Oncogenic and Tumor Suppressor Genes as Top Predictors of Non-Canonical TCA Activity

To gain further insight into model behavior, we applied SHAP analysis to the final XGBoost regressor trained on the stable feature set. SHAP values quantify the marginal contribution of each gene to the predicted activity scores of the non-canonical TCA cycle, providing a ranked list of the most influential features. Also in this case, the model maintained strong predictive performance, with *R*^2^ values consistently above 0.9 across validation folds (*Supplementary Tab. S5*).

We first analyzed SHAP outputs from the model predicting *Cycle Propensity* (*Fig. 4a*). Several top-ranked genes have well-established roles in cancer. *SELENBP1* the top feature, consistent with its function as a tumor suppressor and its frequent downregulation across diverse tumor types [15]. It has also been linked to lipid and glucose metabolism in vivo [16]. *DUSP8*, an inhibitor of the ERK signaling pathway [17]. Its overexpression has been shown to suppress proliferation and migration in colorectal cancer cells [18] *SYT13* another top-ranked gene, promotes proliferation and migration in lung adenocarcinoma, and its silencing reduces metastatic potential in vitro [19]. *DLEU2*, a long non-coding RNA (lncRNA), functions as the host gene of the tumor-suppressive microRNAs *miR-15a* and *miR-16-1*, represses G1 cyclins and inhibits tumor proliferation when overexpressed [20, 21]. *FOXD1* has been shown to drive aerobic glycolysis in pancreatic cancer by upregulating *GLUT1* expression and inhibiting its degradation, thereby promoting cell proliferation, invasion, and metastasis [22]. *HERC5*, has been associated with metabolic reprogramming in non-small cell lung cancer (NSCLC) where reduced expression favors glycolysis, while increased expression promotes oxidative phosphorylation [23].

**Figure 4:**
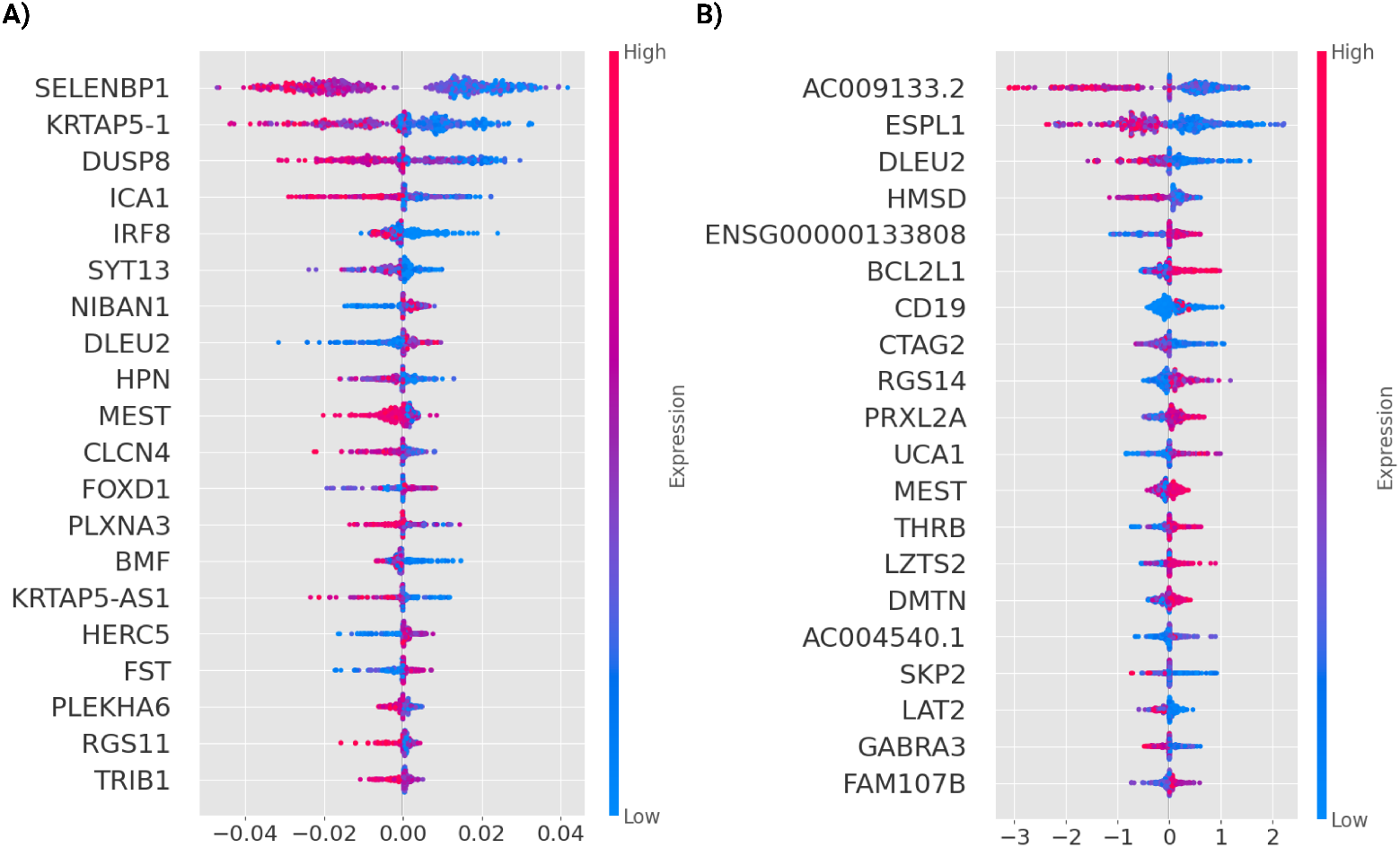
Predictive features of non-canonical TCA cycle activity. **(A)** SHAP summary plot of the top 20 genes contributing most to the prediction of *Cycle Propensity*. Genes are ranked on the y-axis by their overall impact on the model’s output, while the x-axis represents the SHAP value, indicating each gene’s contribution to the deviation from the model’s baseline prediction. Each point corresponds to a cancer cell line, with its color reflecting the gene’s expression level in that cell line. **(B)** SHAP summary plots of the top 20 genes most predictive for the prediction of *Cycle Flux Intensity*.

In the prediction of Cycle Flux Intensity, SHAP analysis revealed a distinct set of influential genes (*Fig. 4b*). *AC009133*.*2*, the top-ranked feature, encodes an antisense lncRNA opposite to *MAZ*, an oncogenic transcription factor implicated in tumor development and metastasis [24]. *ESPL1* which encodes separase, a cysteine protease essential for chromatid separation during mitosis. It is recurrently overexpressed in luminal B breast cancers and is associated with chromosomal instability, loss of tumor suppressor function, and poor clinical outcome [25]. *CTAG2*, a cancer-testis antigen normally restricted to the testis, is aberrantly reactivated in multiple tumor types [26]. In breast cancer, it is required for directional cell migration—a key feature of invasive behavior [27]. *LZTS1*, a tumor suppressor often deleted in epithelial cancers, inhibits invasion and motility, with lower expression correlating with lymph node metastasis in breast cancer [28]. Lastly, *UCA1*, a lncRNA associated with chemoresistance across several cancer types and has been associated with poor therapeutic outcomes [29].

The topological mapping of these SHAP-identified predictors within the PPI network is shown in *Supplementary Fig. S1*. Notably, none of the combined top predictors for *Cycle Propensity* or *Flux Intensity* localized within the large HRAS-centered cluster. By contrast, all four genes forming a smaller subnetwork—*SELENBP1, DMTN, BCL2L1*, and *BMF*—were consistently ranked among the top SHAP-ranked genes. This finding suggests that this cluster may represent a functionally coherent regulatory or effector module associated with non-canonical TCA cycle activity.

Collectively, the top SHAP-identified predictors are functionally implicated in diverse aspects of tumorigenesis, including metabolic reprogramming, proliferation, invasion, and therapeutic resistance.

### 2.5 Genetic Dependencies Associated with Non-Canonical TCA Cycle Propensity

To explore potential associations between non-canonical TCA cycle activity and genetic dependencies in cancer, we correlated DepMap gene dependency scores with *Cycle Propensity*, which reflects the likelihood of cycle engagement across cancer cell lines [30]. Dependency scores represent the impact of gene knockout on cell viability, with more negative values indicating stronger dependency. A score of 0 reflects a non-essential gene, whereas a score of –1 corresponds to the median effect observed for common essential genes.

First, we focused on genes that directly catalyze the reactions of the non-canonical TCA cycle (*Fig. 5a*). As expected, both *ACLY* (*ρ* = −0.26, *p* = 6.43 *×* 10^−7^) and *SLC25A1* (*ρ* = −0.21, *p* = 9.47 *×* 10^−5^) exhibited negative correlations between dependency scores and *Cycle Propensity*, suggesting that cell lines more likely to engage this cycle tend to rely more heavily on these genes. In contrast, *MDH1* showed a weaker but statistically significant positive correlation (*ρ* = 0.15, *p* = 4.55 × 10^−3^), a result that was unexpected and may point to a more nuanced or context-specific role for MDH1 within the non-canonical cycle.

**Figure 5:**
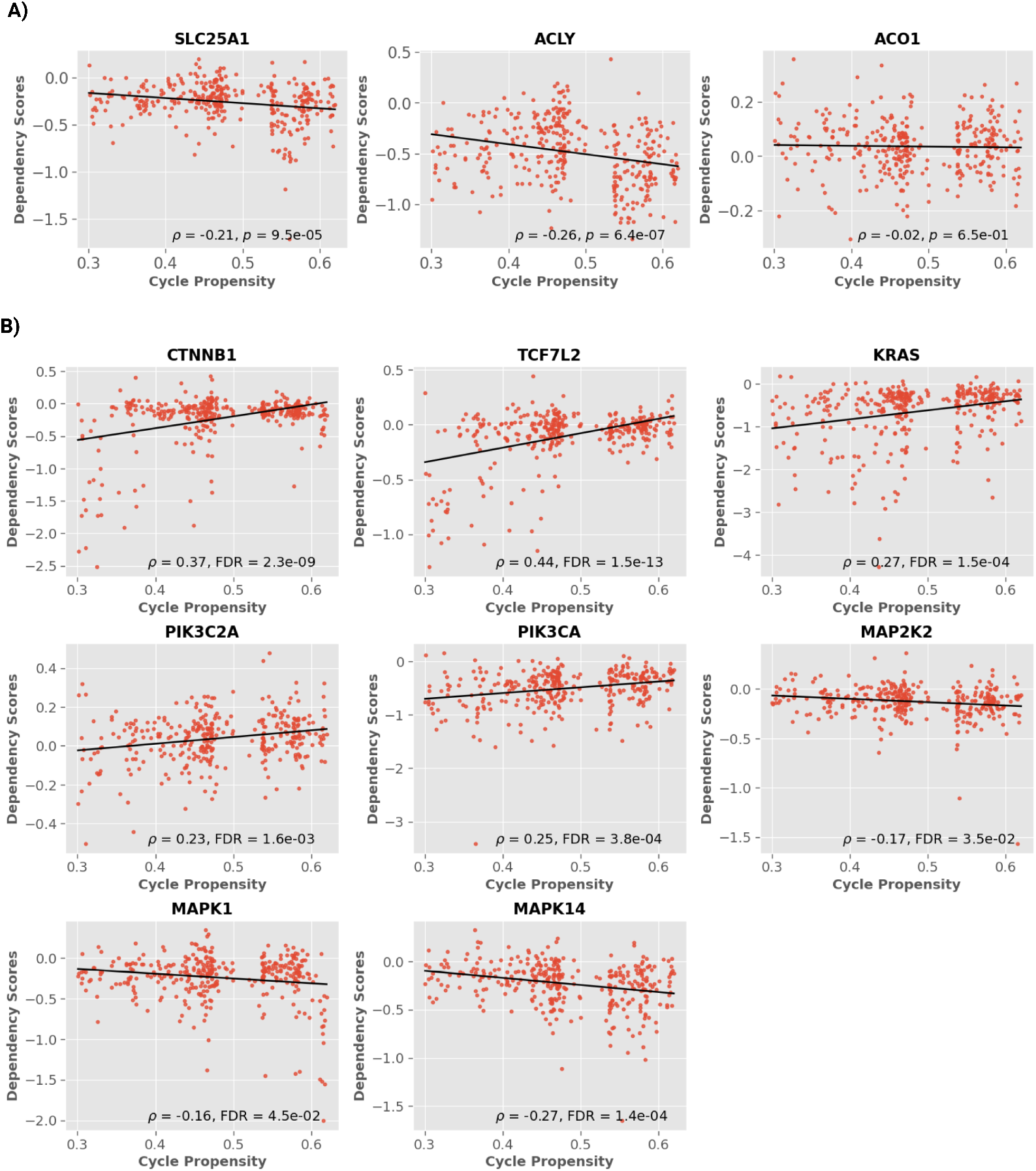
**(A)** Correlation between *Cycle Propensity* and gene dependency scores (DepMap) for enzymes of the Arnold Cycle. Each dot represents a cancer cell line. **(B)** Correlation between *Cycle Propensity* and gene dependency scores for genes in pathways enriched in the over-representation analysis.

To investigate whether engagement of the non-canonical TCA cycle is associated with specific patterns of genetic dependency, we analyzed correlations between Cycle Propensity and gene dependency scores across cancer cell lines (*Fig. 5b*). Notably, many of the top genes whose dependency scores showed the highest correlations with Cycle Propensity are core components of signaling pathways previously enriched in our ORA. These include *TCF7L2* and *CTNNB1* from the canonical Wnt/*β*-catenin pathway; *KRAS, PIK3CA*, and *PIK3C2A*, which are central to the KRAS and PI3K/AKT signaling axes; and multiple members of the MAPK cascade, including *MAPK14, MAP2K2*, and *MAPK1 (ERK2)*. For instance, *TCF7L2* and *CTNNB1* showed strong positive correlations (*ρ* = 0.44, *p* = 1.47 × 10^−13^; *ρ* = 0.37, *FDR* = 2.26 × 10^−9^, respectively), while *KRAS* (*ρ* = 0.27, *FDR* = 1.51 × 10^−4^), *PIK3CA* (*ρ* = 0.25, *FDR* = 3.84 × 10^−4^) and *PIK3C2A* (*ρ* = −0.25, *FDR* = 1.58 × 10^−3^) exhibited moderate correlations. Conversely, multiple components of the MAPK cascade exhibited negative correlations with *Cycle Propensity*, including *MAPK14* (*ρ* = − 0.27, *FDR* = 4.6 × 10^−7^), *MAP2K2* (*ρ* = − 0.17, *FDR* = 0.035), and *MAPK1* (*ρ* = − 0.16, *FDR* = 0.045), suggesting a distinct dependency pattern.

Collectively, these observations suggest that variation in non-canonical TCA cycle propensity across cancer cell lines is associated with selective patterns of genetic dependency involving key oncogenic signaling pathways. The concordance between pathways enriched in the ML-based feature selection (via ORA) and those highlighted by gene dependency correlations reinforces the biological relevance of our computational metrics. This convergence—particularly within the Wnt, KRAS, PI3K/AKT, and MAPK cascades—suggests a potential interplay between metabolic state and oncogenic signaling programs, and provides orthogonal validation that our propensity scores reflect functionally meaningful differences captured by experimental gene perturbation data.

## 3 Discussion

Our study identifies a consistent association between non-canonical TCA cycle activity and altered cancer cell metabolism and transcriptional states. In cell lines with higher Cycle Propensity, cytosolic citrate tends to be increasingly redirected toward *α*KG production via ACO1 and IDH1, rather than being processed by ACLY into acetyl-CoA. This shift reveals that the non-canonical TCA cycle reflects a more intricate metabolic rewiring than previously described by Arnold et al., incorporating not only the canonical ACLY-driven citrate-to-acetyl-CoA route, but also at least one alternative branch that channels citrate toward *α*KG production.

This observation raises the possibility that such metabolic rerouting may influence epigenetic regulation and cell fate decisions through coordinated metabolite availability. We hypothesize that the citrate-to-*α*KG branch may play a regulatory role similar to the canonical citrate-to-acetyl-CoA route, particularly in modulating chromatin dynamics. While acetyl-CoA fuels histone acetylation, *α*KG serves as a cofactor for *α*KG-dependent dioxygenases involved in histone and DNA demethylation. The coordinated availability of these metabolites could synergistically shape gene expression programs associated with cellular differentiation and malignancy. Previous studies have shown that cytosolic IDH1-derived *α*KG modulates histone methylation patterns to control lineage commitment, such as in brown adipocyte differentiation, where *α*KG suppresses adipogenesis by reducing H3K4me3 at key developmental gene promoters [31]. Moreover, mutations in *IDH1* and *IDH2*, recurrently observed in gliomas and acute myeloid leukemias, lead to the neomorphic production of 2-hydroxyglutarate (2HG), a competitive inhibitor of *α*KG-dependent dioxygenases. This aberrant metabolite impairs histone demethylation, resulting in persistent repression of differentiation-associated genes and maintenance of a progenitor-like transcriptional state [32].

This rerouting of cytosolic citrate may not occur in isolation, but rather as part of a broader metabolic reprogramming. One of the most well-characterized examples of such reprogramming is the Warburg effect. Indeed, our findings reveal that increased engagement of the non-canonical TCA cycle is closely associated with a Warburg-like metabolic phenotype. In particular, earlier research has demonstrated that that ACLY, promotes glycolytic metabolism and lactate production in cancer cells. For example, its overexpression in breast cancer enhances glycolytic enzyme expression, glucose uptake, and lactate secretion, facilitating tumor progression via epithelial-to-mesenchymal transition (EMT) [33]. Similarly, in glioblastoma, ACLY accumulation in pseudopodia sustains migration and preserves glycolytic flux under mitochondrial inhibition [34]. In light of its proposed role in chromatin remodeling, we next asked whether the metabolic activity of the non-canonical TCA cycle is coupled to specific transcriptional programs. To address this, we performed Over-Representation Analysis on genes identified through machine learning-based feature selection. This analysis uncovered a robust transcriptional signature associated with non-canonical TCA cycle activity, enriched for multiple hallmarks of malignancy—including invasion, stemness, and oncogenic signaling pathways—with strong concordance to prior experimental studies. Interestingly, we also observed enrichment for angiogenesis-related processes, a novel association that, to our knowledge, has not been directly linked to enzymes of the non-canonical TCA cycle. This finding highlights a potentially unexplored connection between metabolic rewiring and vascular regulation, warranting further experimental validation.

Specifically, ACLY has been shown to enhance metastatic behavior in colon cancer by stabilizing *β*-catenin and increasing its transcriptional activity, thereby facilitating EMT and the induction of pro-metastatic genes [35]. A similar pro-invasive role for ACLY has been reported in hepatocellular carcinoma (HCC), further highlighting its contribution to metastatic competence [36]. Beyond its role in invasion, ACLY and related metabolic enzymes have been functionally linked to several oncogenic pathways enriched in our analysis. ACLY is a known downstream target of the PI3K/AKT pathway and is activated through direct AKT-mediated phosphorylation [37]. The observed enrichment for Wnt signaling is also consistent with reports connecting ACLY to *β*-catenin stabilization [35]. In addition, the prominence of KRAS and ERK1/2 signaling cascades may reflect broader layers of metabolic–signaling crosstalk. It has been proposed that ACLY-driven glycolytic activation elevates fructose-1,6-bisphosphate levels, which in turn promote RAS activation via Sos1, ultimately reinforcing MEK/ERK and PI3K/AKT signaling loops [38].

We also observed enrichment for terms associated with stemness and developmental plasticity. These results echo previous reports implicating ACLY in the maintenance of cancer stem cell (CSC) features. Specifically, knockdown of ACLY has been shown to suppress self-renewal capacity and tumorsphere formation in models of NSCLC, HCC, and breast cancer [39, 36]. Additionally, the mitochondrial citrate transporter SLC25A1 has been implicated in CSC regulation, with its overexpression in NSCLC models promoting self-renewal and enhancing the expression of canonical stemness markers [40]. Recent work in liver cancer stem cells extends this connection, showing that SLC25A1 is essential for mitochondrial function and self-renewal capacity. [41]. The overlap between signaling pathways enriched in ORA and those identified through gene dependency correlations reinforces the robustness and biological relevance of our cycle activity quantification metrics, and provides support for the validity of our computational framework.

The genes identified through our machine learning approach constitute valuable candidates for targeted validation experiments, despite the inherent limitation of our methodology in distinguishing upstream regulators from downstream effectors. One potential discrimination strategy would involve assessing histone acetylation at promoters of selected genes, given that acetyl-CoA production through ACLY directly contributes to histone acetylation. Unfortunately, this specific type of epigenetic data is currently unavailable in the CCLE dataset.

Finally, although our analysis centers on the non-canonical TCA cycle, the modular framework developed here is broadly applicable to the study of other metabolic pathways. However, it is essential to emphasize that the performance of our pipeline is highly dependent on the accuracy and completeness of the underlying metabolic model reconstruction. Refinement of these models will be essential to ensure that the derived outputs reflect biologically meaningful and context-specific insights.

## 4 Methods

### 4.1 Overview of the Computational Framework

We adopted a two-step computational strategy to dissect Arnold Cycle activity across cancer cell lines (*Fig. 1c*).

In the first module, we used constraint-based metabolic modeling to infer the activity of the Arnold Cycle from gene expression data. Specifically, we extracted transcriptomic profiles from 513 cancer cell lines cultured under standardized conditions and reconstructed a cell-specific metabolic model for each line based on the ENGRO2.2 core network. Reaction constraints were individualized by computing Reaction Activity Scores (RAS) using Gene–Protein–Reaction (GPR) rules, and these scores were used to rescale flux boundaries to reflect the transcriptional context of each cell line [42]. To comprehensively explore the space of feasible metabolic states, we performed flux sampling on each transcriptome-constrained model using a corner-based sampling algorithm that explores the solution space without relying on a predefined objective function. This allowed us to sample steady-state flux configurations consistent with mass balance and expression-derived constraints, without assuming predefined cellular objectives. From these sampled flux distributions, we derived two complementary metrics to quantify Arnold Cycle activity: *Cycle Propensity*, defined as the fraction of flux samples in which the three reactions (*SLC25A1, ACLY, MDH1*) of the Arnold Cycle are simultaneously active *Fig. 1b*); and *Cycle Flux Intensity*, which captures the average flux through the bottleneck reaction when the cycle is active.

In the second module, these *in silico*–derived metrics were used as training labels for supervised machine learning models. To predict Cycle Propensity and Cycle Flux Intensity from transcriptional features beyond the metabolic network, we trained regressors using expression data from non-metabolic genes. After model training, we performed feature selection to identify robust predictors of Arnold Cycle activity, thereby uncovering broader transcriptional programs associated with its engagement.

### 4.2 Data collection and pre-processing

The gene expression data were obtained from the Cancer Cell Line Encyclopedia (CCLE) [43], including RNA-seq profiles of 1,019 human cancer cell lines, normalized to Transcripts Per Million (TPM). To minimize experimental variability arising from heterogeneous culture conditions, we restricted the dataset to 513 cell lines cultured exclusively in RPMI-1640 medium—the most prevalent condition in the dataset. This homogenization step minimized confounding variables during subsequent metabolic model reconstruction.

Gene expression values were log-transformed using the `log1p` function and imputed using MAGIC (Markov Affinity-based Graph Imputation of Cells) with default parameters (k=5, t=3) [44]. Although denoising is more commonly applied to single-cell RNA-sequencing data [45]—where sparsity arises from dropout events—we found that applying it to our bulk transcriptomic profiles substantially improved downstream analyses. In particular, it yielded higher silhouette scores when clustering cell lines based on simulated flux distributions, and also led to significantly better alignment with transcriptome-based clusters defined by metabolic gene expression. In the absence of denoising, the two clustering solutions showed low concordance. These preprocessed data were used directly for cell-specific metabolic model reconstruction.

For downstream machine learning analyses, we further filtered the expression matrix using a four-step procedure. First, genes with low cross-sample variability (variance < 0.1) were excluded due to limited predictive utility. Second, genes expressed in fewer than 30 cell lines were removed to eliminate tissue-specific genes and enhance generalizability across tumour types. Third, all annotated pseudogenes were filtered out using Ensembl biotype classifications, as they are unlikely to be functionally relevant. Finally, genes encoding metabolic enzymes were systematically excluded to avoid circularity in downstream analyses and to focus the model on non-metabolic regulators of non-canonical TCA cycle activity. The list of metabolic genes was curated from the Human Protein Atlas.

The resulting expression matrix was standardized using `StandardScaler` to harmonize feature scales and facilitate convergence in regularized learning algorithms.

### 4.3 Metabolic Model

Cell line–specific models were built based on the recently published ENGRO2 core network model, a curated reconstruction of human central carbon metabolism [2]. We chose a network of reduced complexity over genome-scale reconstructions because of its greater computational efficiency and interpretability. While genome-scale models provide broader metabolic coverage, their inclusion of numerous alternative pathways can obscure specific metabolic activities in flux simulations. In line with this, preliminary flux sampling experiments using the Recon3D [46] model revealed that both canonical and non-canonical TCA cycles were poorly represented. These results support the use of a core model for our analysis.

Compared to the original ENGRO2 network [2], we previously added the reactions related to the galactose and cholesterol uptake. Moreover, we revised the GPR rules related to amino acid uptake[45]. In particular, we incorporate the information about the gene expression of all relevant amino acid transporters, extending the GPRs of the uptake reactions present in the original

ENGRO2 model. Specifically, for each amino acid, we used the OR logical operator to merge the GPRs of all known uptake reactions involving that amino acid, as identified in the Recon3D metabolic network [46]. Finally, to adapt the model for the current analysis, we incorporated literature-supported alternative pathways for cytosolic acetyl-CoA production, including two endogenous acetate generation mechanisms, to better reflect the metabolic flexibility relevant to this study [47, 48, 49]. This extended version, hereafter referred to as ENGRO2.2, comprises 395 metabolites, 469 reactions, and 498 genes. Among the 469 reactions, 351 are associated with GPR rules, enabling robust integration of transcriptomic data into the model.

The biomass pseudo-reaction of ENGRO2 corresponds with the biomass reaction of the Recon3D model, in terms of the set of metabolites considered and their corresponding stoichiometric coefficients.

### 4.4 Cell Line–Specific Model Reconstruction

#### 4.4.1 Reaction Activity Score Calculation

RAS were calculated for each reaction across all cell line models by GPR rules based on mRNA abundance data, as described previously [42]. Logical expressions were resolved such that the minimum transcript level was used for AND logic, while the sum was used for OR logic. The standard precedence (AND before OR) was applied when both operators appeared in a GPR rule. Following RAS computation for all reactions in the metabolic model, a cell line–by–reaction matrix of RAS values was generated.

#### 4.4.2 Flux Variability Analysis

To characterize the feasible flux space of the metabolic network, defined by mass balance constraints and medium composition, we employed Flux Variability Analysis (FVA) [50, 51, 52]. This constraint-based approach computes the minimum and maximum possible flux 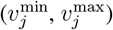 for each reaction *j* (*j* = 1, …, *R*) subject to the defined constraints.

For each reaction *j*, FVA solves the optimization problem:

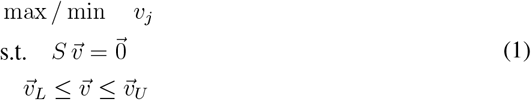

where *S* is the stoichiometric matrix, 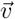 the flux vector, 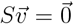 enforces steady-state mass balance, and 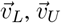 represent flux bounds imposed by biological constraints and the RPMI-1640 medium.

#### 4.4.3 Cell Line–Specific Flux Constraints

Based on the computed RAS and FVA results, cell line–specific flux constraints were established using an approach adapted from [53]. For each reaction *j* (*j* = 1, …, *R*) and cell line *c* (*c* = 1, …, *C*), the flux bounds were computed as follows:

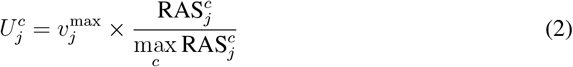

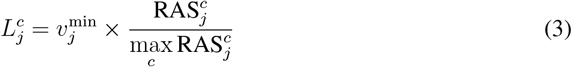

where 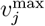 and 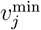 are the upper and lower bounds obtained from Flux Variability Analysis (Section 4.4.2), 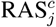 denotes the Reaction Activity Score for reaction *j* in cell line *c* (Section 4.4.1), and 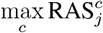 is the maximum RAS value for reaction *j* across all cell lines.

Additionally, the lower bound for the biomass reaction was set to 0.01 to enforce a minimal proliferation, reflecting the assumption that each cell line sustains a basal level of growth sufficient to preserve population viability. Therefore, each cell cine–specific model has specific constraints derived from the transcriptomics and defined in Eq. 3 and 2

### 4.5 Flux Sampling

To explore the feasible flux space of each cell-specific metabolic model without assuming a predefined objective, we employed flux sampling. This method generates multiple flux distributions that satisfy steady-state and constraint-based conditions, offering a probabilistic view of metabolic activity beyond the scope of traditional objective-driven approaches like FBA.

In this study, we employed the Corner-based Sampling (CBS) algorithm to sample the vertices of the feasible flux space by iteratively optimizing randomly weighted objective functions [54, 55]. At each iteration, weights were assigned to reactions by sampling from a uniform distribution in the range [− 1, 1], and the objective was randomly selected for maximization or minimization. To account for differences in reaction scales, each weight was normalized by the corresponding maximum flux value obtained from FVA. Further implementation details are available in [55]. For each cell-specific model, we generated 10,000 flux samples.

### 4.6 Non-Canonical TCA Activity Metrics

To quantify the activity of the non-canonical TCA cycle across cell lines, we defined two complementary metrics: *Cycle Propensity* and *Cycle Flux Intensity*. Both were computed independently for each cell-specific metabolic model, indexed by *c*.

***Cycle Propensity*** captures the likelihood with which a cell line activates the non-canonical cycle. This was defined as the fraction of sampled flux distributions in which all three hallmark reactions—citrate export via SLC25A1, cleavage by ACLY, and malate regeneration via MDH1—exceed a minimal flux threshold (*ε* = 10^−6^) and operate in the expected physiological direction.

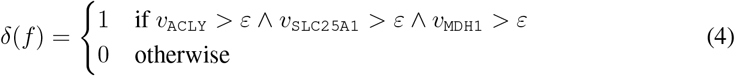

Here, *δ*(*f*) is a binary indicator function assigning 1 to active flux samples and 0 otherwise; *ν*_ACLY_, *ν*_SLC25A1_, *ν*_MDH1_ denote the fluxes through the corresponding reactions in sample *f*.

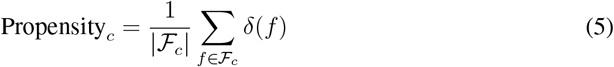

In this expression, *ℱ*_*c*_ is the set of all flux samples generated for cell line *c*.

***Cycle Flux Intensity*** measures the average operational strength of the cycle, conditional on activation. For each reaction *r ∈ R* = {`ACLY`, `SLC25A1`, `MDH1`}, we computed the mean flux across the subset of active samples. The final intensity value was defined as the minimum of these means, representing the rate-limiting step of the cycle:

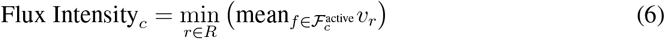

where 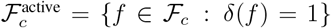. This formulation ensures that the metric reflects the bottleneck flux among the core reactions, thus offering a conservative estimate of cycle throughput in cell line c.

### 4.7 Machine Learning Framework

To identify genes whose expression is associated with non-canonical TCA cycle activity across different cancer cell lines, we developed a supervised regression pipeline to predict two activity metrics—*Cycle Propensity* and *Cycle Flux Intensity*, as described in Section 4.6. The input matrix *X* consisted of preprocessed gene expression data, while the output matrix *y* contained the two corresponding activity metrics for each cell line. The mean and standard deviation of these two target variables are reported in **Supplementary Tab. S2**

To capture both linear and non-linear associations between gene expression and metabolic phenotypes, we adopted two complementary feature selection strategies: *ElasticNet*, a linear model promoting variable selection through *ℓ*_1_/*ℓ*_2_ regularization, and *Random Forest feature importance*, a non-parametric ensemble method capable of modeling complex, higher-order interactions among features.

#### 4.7.1 ElasticNet Feature Selection

We applied ElasticNet regression, a regularized linear model combining *ℓ*_1_ and *ℓ*_2_ penalties, to model associations between gene expression and non-canonical TCA cycle activity [56]. The model was implemented using `MultiOutputRegressor`, which independently fits one ElasticNet model for each target variable.

Model training and feature selection were embedded in a nested cross-validation framework with 10 outer folds and 5 inner folds. The outer loop assessed model generalizability and feature stability, while the inner loop was dedicated to hyperparameter optimization. We employed Bayesian search using the Tree-structured Parzen Estimator (TPE) implemented in the `Hyperopt` library [57]. The search space included regularization strength, *ℓ*_1_/*ℓ*_2_ mixing ratio, maximum number of iterations, and convergence tolerance. The inner loop objective was to minimize the mean squared error across validation folds. Performance metrics for the ElasticNet models are reported in **Supplementary Tab. S3**

Due to its *ℓ*_1_ component, ElasticNet yields sparse solutions by setting many regression coefficients to zero. For each outer fold, we retained genes with non-zero coefficients in at least one of the two target-specific models. Binary selection masks were stored for all folds, enabling the computation of per-gene selection frequencies and assessment of feature stability.

#### 4.7.2 Non-Linear Feature Selection with Random Forests

We applied Random Forest regression to model gene expression–based predictions of non-canonical TCA cycle activity. [58]. As in the ElasticNet framework, we used `MultiOutputRegressor` to fit separate Random Forest models for each output variable.

Model selection and feature assessment were performed using the same nested cross-validation framework (10 outer folds, 5 inner folds). Hyperparameter tuning was carried out in the inner loop using the Tree-structured Parzen Estimator (TPE) in `Hyperopt`, minimizing the mean squared error across validation sets. Tuned parameters included the number of estimators, maximum tree depth, minimum number of samples per split and per leaf, and the maximum fraction of features considered at each split. Performance metrics for the Random Forest models are reported in **Supplementary Tab. S4**

After training each final model, average feature importance across trees was computed. Unlike ElasticNet, Random Forest inherently assigns non-zero importance to all features; therefore, we applied a cumulative importance thresholding strategy to identify the most informative genes. For each fold, features were ranked in descending order of importance, and only those contributing to the top 85% of the total cumulative importance were retained.

#### 4.7.3 Stable Feature Selection

To ensure the robustness of selected features, we quantified their stability across outer cross-validation folds. For each predictive model (ElasticNet and Random Forest), we generated binary selection masks indicating whether each gene was selected in a given fold.

For gene *i*, a stability score *s*_*i*_ ∈ [0, 1] was computed as the proportion of outer folds in which the gene was selected:

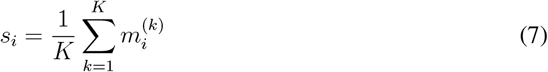

where *K* = 10 is the number of outer folds and 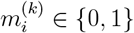 indicates whether gene *i* was selected in fold *k*. Genes with *s*_*i*_ ≥ 0.8 were considered stable, reflecting high recurrence across folds.

Applying this criterion, we identified 94 stable features from ElasticNet, 226 from Random Forest, and 6 genes consistently selected by both methods. complete feature list is reported in **Supplementary Tab. S1**.

#### 4.7.4 Gene-Level Interpretation via XGBoost and SHAP

To quantify the predictive contribution of individual genes, we trained eXtreme Gradient Boosting (XGBoost) models using the stable feature sets identified in the previous step [59]. XGBoost was wrapped within a `MultiOutputRegressor` framework, fitting one model for each activity metric.

Model performance and generalizability were assessed using 10-fold nested cross-validation. Hyperparameters were optimized in the inner loop via Bayesian search using the Tree-structured Parzen Estimator (TPE) implemented in `Hyperopt`. The search space included the number of estimators, maximum tree depth, learning rate, minimum child weight, gamma, subsample ratio, column sample ratio, and both *ℓ*_1_ and *ℓ*_2_ regularization terms. Performance metrics for the XGBoos models are reported in **Supplementary Tab. S5**

To interpret model predictions, we applied SHAP [60]. For each outer fold, SHAP values were computed on the test set using `TreeExplainer`, separately for the two fitted estimators. SHAP values across all outer test sets were concatenated, and gene-level importance was summarized as the mean absolute SHAP value, yielding a global ranking of predictive relevance.

### 4.8 Protein–Protein Interaction Network Analysis

We constructed a protein–protein interaction (PPI) network using the STRING database *ver. 12*.*0* [61], integrating experimental evidence, curated database entries, and co-expression data, with a minimum interaction confidence score of 0.4. Network clustering was performed using the Markov Clustering Algorithm with an inflation parameter of 1.2, resulting in 23 distinct clusters, including one major module and several smaller groups.

### 4.9 Over Representation Analysis

Over-Representation Analysis was conducted on the final set of 314 selected genes. Enrichment was tested across four major gene set collections: Gene Ontology Biological Processes, KEGG, Reactome, and MSigDB Hallmark gene sets.

The background gene set included 32,555 expressed transcripts, excluding pseudogenes and canonical metabolic genes. Enrichment significance was assessed using a hypergeometric test with Benjamini–Hochberg correction for multiple testing, and only terms with an FDR below were considered significant.

ORA was carried out using the online tool `g:Profiler` [62], *ver. e113_eg59_p19_f6a03c19*, with electronic GO annotations disabled to avoid low-confidence associations. To reduce redundancy among enriched GO terms, results were post-processed using `REViGO` *ver. 1*.*8*.*1* [63]. Redundancy filtering was applied using the “Small (0.5)” setting for semantic similarity.

### 4.10 Gene Dependency Correlation Analysis

We correlated Cycle Propensity scores with genome-wide gene dependency scores from the Cancer Dependency Map project *ver. 24Q2* [30]. We used CERES gene effect scores.

Pearson correlation coefficients were computed between Cycle Propensity and gene dependency scores across all cancer cell lines for which both datasets were available. Statistical significance was adjusted using the Benjamini–Hochberg, and only genes with *FDR* < 0.05 were retained for downstream analysis.

## Data and Code Availability

The transcriptomic data and gene dependency scores used in this study are publicly available from the Cancer Cell Line Encyclopedia and DepMap.

All code used for data preprocessing, flux sampling, and the machine learning pipeline—along with processed datasets and a reproducible workflow—is available at: https://github.com/CompBtBs/noncanonical-tca-cancer-analysis

## CRediT authorship contribution statement

**Lihao Lin**: Conceptualization; Investigation; Methodology; Software; Visualization; Writing – original draft; Writing – review & editing. **Francesco Lapi**: Investigation; Methodology; Software. **Bruno G. Galuzzi**: Methodology; Software. **Marco Vanoni**: Conceptualization; Funding acquisition; Writing – review & editing. **Lilia Alberghina**: Conceptualization; Funding acquisition. **Chiara Damiani**: Conceptualization; Methodology; Supervision; Funding acquisition; Writing – review & editing.

## References

[1] Paige K Arnold, Benjamin T Jackson, Katrina I Paras, Julia S Brunner, Madeleine L Hart, Oliver J Newsom, Sydney P Alibeckoff, Jennifer Endress, Esther Drill, Lucas B Sullivan, et al. A non-canonical tricarboxylic acid cycle underlies cellular identity. Nature, 603(7901):477–481, 2022.

[2] Marzia Di Filippo, Dario Pescini, Bruno Giovanni Galuzzi, Marcella Bonanomi, Daniela Gaglio, Eleonora Mangano, Clarissa Consolandi, Lilia Alberghina, Marco Vanoni, and Chiara Damiani. Integrate: Model-based multi-omics data integration to characterize multi-level metabolic regulation. PLoS computational biology, 18(2):e1009337, 2022.

[3] Edward J O’Brien, Jonathan M Monk, and Bernhard O Palsson. Using genome-scale models to predict biological capabilities. Cell, 161(5):971–987, 2015.

[4] Daniel Machado and Markus Herrgård. Systematic evaluation of methods for integration of transcriptomic data into constraint-based models of metabolism. PLoS computational biology, 10(4):e1003580, 2014.

[5] David A Guertin and Kathryn E Wellen. Acetyl-coa metabolism in cancer. Nature Reviews Cancer, 23(3):156– 172, 2023.

[6] Douglas C Wallace. Mitochondria and cancer. Nature Reviews Cancer, 12(10):685–698, 2012.

[7] Otto Warburg. The metabolism of carcinoma cells. The Journal of Cancer Research, 9(1):148–163, 1925.

[8] Matthew G Vander Heiden, Lewis C Cantley, and Craig B Thompson. Understanding the warburg effect: the metabolic requirements of cell proliferation. science, 324(5930):1029–1033, 2009.

[9] Maria V Liberti and Jason W Locasale. The warburg effect: how does it benefit cancer cells? Trends in biochemical sciences, 41(3):211–218, 2016.

[10] Lilia Alberghina. The warburg effect explained: integration of enhanced glycolysis with heterogeneous mitochondria to promote cancer cell proliferation. International Journal of Molecular Sciences, 24(21):15787, 2023.

[11] Douglas Hanahan and Robert A Weinberg. Hallmarks of cancer: the next generation. cell, 144(5):646–674, 2011.

[12] Yan-Jun Guo, Wei-Wei Pan, Sheng-Bing Liu, Zhong-Fei Shen, Ying Xu, and Ling-Ling Hu. Erk/mapk signalling pathway and tumorigenesis. Experimental and therapeutic medicine, 19(3):1997–2007, 2020.

[13] David Tai, Keith Wells, John Arcaroli, Chad Vanderbilt, Dara L Aisner, Wells A Messersmith, and Christopher H Lieu. Targeting the wnt signaling pathway in cancer therapeutics. The oncologist, 20(10):1189–1198, 2015.

[14] Saeed Noorolyai, Neda Shajari, Elham Baghbani, Sanam Sadreddini, and Behzad Baradaran. The relation between pi3k/akt signalling pathway and cancer. Gene, 698:120–128, 2019.

[15] Mostafa Elhodaky and Alan M Diamond. Selenium-binding protein 1 in human health and disease. International journal of molecular sciences, 19(11):3437, 2018.

[16] Qi Ying, Emmanuel Ansong, Alan M Diamond, Zhaoxin Lu, Wancai Yang, and Xiaomei Bie. Quantitative proteomic analysis reveals that anti-cancer effects of selenium-binding protein 1 in vivo are associated with metabolic pathways. PloS one, 10(5):e0126285, 2015.

[17] Tao Ding, Ya Zhou, Runying Long, Chao Chen, Juanjuan Zhao, Panpan Cui, Mengmeng Guo, Guiyou Liang, and Lin Xu. Dusp8 phosphatase: structure, functions, expression regulation and the role in human diseases. Cell & Bioscience, 9(1):70, 2019.

[18] Tao Ding, Panpan Cui, Ya Zhou, Chao Chen, Juanjuan Zhao, Hairong Wang, Mengmeng Guo, Zhixu He, and Lin Xu. Antisense oligonucleotides against mir-21 inhibit the growth and metastasis of colorectal carcinoma via the dusp8 pathway. Molecular Therapy-Nucleic Acids, 13:244–255, 2018.

[19] Liyan Zhang, Bijun Fan, Yu Zheng, Yueyan Lou, Yongqi Cui, Ke Wang, Tiancheng Zhang, and Xiaoming Tan. Identification syt13 as a novel biomarker in lung adenocarcinoma. Journal of Cellular Biochemistry, 121(2):963–973, 2020.

[20] Ulf Klein, Marie Lia, Marta Crespo, Rachael Siegel, Qiong Shen, Tongwei Mo, Alberto Ambesi-Impiombato, Andrea Califano, Anna Migliazza, Govind Bhagat, et al. The dleu2/mir-15a/16-1 cluster controls b cell proliferation and its deletion leads to chronic lymphocytic leukemia. Cancer cell, 17(1):28–40, 2010.

[21] Mikael Lerner, Masako Harada, Jakob Lovén, Juan Castro, Zadie Davis, David Oscier, Marie Henriksson, Olle Sangfelt, Dan Grandér, and Martin M Corcoran. Dleu2, frequently deleted in malignancy, functions as a critical host gene of the cell cycle inhibitory micrornas mir-15a and mir-16-1. Experimental cell research, 315(17):2941–2952, 2009.

[22] Kun Cai, Shiyu Chen, Changhao Zhu, Lin Li, Chao Yu, Zhiwei He, and Chengyi Sun. Foxd1 facilitates pancreatic cancer cell proliferation, invasion, and metastasis by regulating glut1-mediated aerobic glycolysis. Cell Death & Disease, 13(9):765, 2022.

[23] Svenja Schneegans, Jana Löptien Angelika Mojzisch, Desirée Loreth, Oliver Kretz, Christoph Raschdorf, Annkathrin Hanssen, Antonia Gocke, Bente Siebels, Karthikeyan Gunasekaran, et al. Herc5 downregulation in non-small cell lung cancer is associated with altered energy metabolism and metastasis. Journal of Experimental & Clinical Cancer Research, 43(1):110, 2024.

[24] Chuanjun Zheng, Hongmei Wu, Song Jin, D. Li, Shengkui Tan, and Xiaonian Zhu. Roles of myc-associated zinc finger protein in malignant tumors. Asia-Pacific Journal of Clinical Oncology, 18(6):506–514, 2022.

[25] Pascal Finetti, Arnaud Guille, José Adelaide Daniel Birnbaum, Max Chaffanet, and François Bertucci. Espl1 is a candidate oncogene of luminal b breast cancers. Breast cancer research and treatment, 147:51–59, 2014.

[26] Zane A Gibbs and Angelique W Whitehurst. Emerging contributions of cancer/testis antigens to neoplastic behaviors. Trends in cancer, 4(10):701–712, 2018.

[27] Erin A Maine, Jill M Westcott, Amanda M Prechtl, Tuyen T Dang, Angelique W Whitehurst, and Gray W Pearson. The cancer-testis antigens spanx-a/c/d and ctag2 promote breast cancer invasion. Oncotarget, 7(12):14708, 2016.

[28] Xin-Xin Wang, Bing-Bing Liu, Xiao Wu, Dan Su, Zhengmao Zhu, and Li Fu. Loss of leucine zipper putative tumor suppressor 1 (lzts1) expression contributes to lymph node metastasis of breast invasive micropapillary carcinoma. Pathology & Oncology Research, 21:1021–1026, 2015.

[29] Haohao Wang, Zhonghai Guan, Kuifeng He, Jiong Qian, Jiang Cao, and Lisong Teng. Lncrna uca1 in anti-cancer drug resistance. Oncotarget, 8(38):64638, 2017.

[30] Aviad Tsherniak, Francisca Vazquez, Phil G Montgomery, Barbara A Weir, Gregory Kryukov, Glenn S Cowley, Stanley Gill, William F Harrington, Sasha Pantel, John M Krill-Burger, et al. Defining a cancer dependency map. Cell, 170(3):564–576, 2017.

[31] Hyun Sup Kang, Jae Ho Lee, Kyoung-Jin Oh, Eun Woo Lee, Baek Soo Han, Kun-Young Park, Jae Myoung Suh, Jeong-Ki Min, Seung-Wook Chi, Sang Chul Lee, et al. Idh1-dependent *α*-kg regulates brown fat differentiation and function by modulating histone methylation. Metabolism, 105:154173, 2020.

[32] Chao Lu, Patrick S Ward, Gurpreet S Kapoor, Dan Rohle, Sevin Turcan, Omar Abdel-Wahab, Christopher R Edwards, Raya Khanin, Maria E Figueroa, Ari Melnick, et al. Idh mutation impairs histone demethylation and results in a block to cell differentiation. Nature, 483(7390):474–478, 2012.

[33] Yu Lu, Liping Tian, Chengcheng Peng, Jienan Kong, Pengpeng Xiao, and Nan Li. Acly-induced reprogramming of glycolytic metabolism plays an important role in the progression of breast cancer: Role of acly in breast cancer. Acta Biochimica et Biophysica Sinica, 55(5):878, 2023.

[34] Marie E Beckner, Wendy Fellows-Mayle, Zhe Zhang, Naomi R Agostino, Jeffrey A Kant, Billy W Day, and Ian F Pollack. Identification of atp citrate lyase as a positive regulator of glycolytic function in glioblastomas. International Journal of Cancer, 126(10):2282–2295, 2010.

[35] Jun Wen, Xuejie Min, Mengqin Shen, Qian Hua, Yuan Han, Li Zhao, Liu Liu, Gang Huang, Jianjun Liu, and Xiaoping Zhao. Acly facilitates colon cancer cell metastasis by ctnnb1. Journal of Experimental & Clinical Cancer Research, 38:1–12, 2019.

[36] Qin Han, Ci-An Chen, Wen Yang, Dong Liang, Hong-Wei Lv, Gui-Shuai Lv, Qian-Ni Zong, and Hong-Yang Wang. Atp-citrate lyase regulates stemness and metastasis in hepatocellular carcinoma via the wnt/*β*-catenin signaling pathway. Hepatobiliary & Pancreatic Diseases International, 20(3):251–261, 2021.

[37] Gerta Hoxhaj and Brendan D Manning. The pi3k–akt network at the interface of oncogenic signalling and cancer metabolism. Nature Reviews Cancer, 20(2):74–88, 2020.

[38] Philippe Icard, Zherui Wu, Ludovic Fournel, Antoine Coquerel, Hubert Lincet, and Marco Alifano. Atp citrate lyase: A central metabolic enzyme in cancer. Cancer letters, 471:125–134, 2020.

[39] JI Hanai, N Doro, P Seth, and VP Sukhatme. Atp citrate lyase knockdown impacts cancer stem cells in vitro. Cell death & disease, 4(6):e696–e696, 2013.

[40] Harvey R Fernandez, Shreyas M Gadre, Mingjun Tan, Garrett T Graham, Rami Mosaoa, Martin S Ongkeko, Kyu Ah Kim, Rebecca B Riggins, Erika Parasido, Iacopo Petrini, et al. The mitochondrial citrate carrier, slc25a1, drives stemness and therapy resistance in non-small cell lung cancer. Cell Death & Differentiation, 25(7):1239–1258, 2018.

[41] Zhichun Zhang, Yuan Qiao, Qiuyue Sun, Liang Peng, and Lichao Sun. A novel slc25a1 inhibitor, parthenolide, suppresses the growth and stemness of liver cancer stem cells with metabolic vulnerability. Cell Death Discovery, 9(1):350, 2023.

[42] Alex Graudenzi, Davide Maspero, Marzia Di Filippo, Marco Gnugnoli, Claudio Isella, Giancarlo Mauri, Enzo Medico, Marco Antoniotti, and Chiara Damiani. Integration of transcriptomic data and metabolic networks in cancer samples reveals highly significant prognostic power. Journal of biomedical informatics, 87:37–49, 2018.

[43] Mahmoud Ghandi, Franklin W Huang, Judit Jané-Valbuena, Gregory V Kryukov, Christopher C Lo, E Robert McDonald III, Jordi Barretina, Ellen T Gelfand, Craig M Bielski, Haoxin Li, et al. Next-generation characterization of the cancer cell line encyclopedia. Nature, 569(7757):503–508, 2019.

[44] David Van Dijk, Roshan Sharma, Juozas Nainys, Kristina Yim, Pooja Kathail, Ambrose J Carr, Cassandra Burdziak, Kevin R Moon, Christine L Chaffer, Diwakar Pattabiraman, et al. Recovering gene interactions from single-cell data using data diffusion. Cell, 174(3):716–729, 2018.

[45] Bruno G Galuzzi, Stefano Izzo, Fabio Giampaolo, Salvatore Cuomo, Marco E Vanoni, Lilia Alberghina, Chiara Damiani, and Francesco Piccialli. Coupling constrained-based flux sampling and clustering to tackle cancer metabolic heterogeneity. In 2023 31st Euromicro International Conference on Parallel, Distributed and Network-Based Processing (PDP), pages 185–192. IEEE, 2023.

[46] Elizabeth Brunk, Swagatika Sahoo, Daniel C Zielinski, Ali Altunkaya, Andreas Dräger, Nathan Mih, Francesco Gatto, Avlant Nilsson, German Andres Preciat Gonzalez, Maike Kathrin Aurich, et al. Recon3d enables a three-dimensional view of gene variation in human metabolism. Nature biotechnology, 36(3):272–281, 2018.

[47] Steven Zhao, AnnMarie Torres, Ryan A Henry, Sophie Trefely, Martina Wallace, Joyce V Lee, Alessandro Carrer, Arjun Sengupta, Sydney L Campbell, Yin-Ming Kuo, et al. Atp-citrate lyase controls a glucose-to-acetate metabolic switch. Cell reports, 17(4):1037–1052, 2016.

[48] Xiaojing Liu, Daniel E Cooper, Ahmad A Cluntun, Marc O Warmoes, Steven Zhao, Michael A Reid, Juan Liu, Peder J Lund, Mariana Lopes, Benjamin A Garcia, et al. Acetate production from glucose and coupling to mitochondrial metabolism in mammals. Cell, 175(2):502–513, 2018.

[49] Luke T Izzo, Sophie Trefely, Christina Demetriadou, Jack M Drummond, Takuya Mizukami, Nina Kuprasertkul, Aimee T Farria, Phuong TT Nguyen, Nivitha Murali, Lauren Reich, et al. Acetylcarnitine shuttling links mitochondrial metabolism to histone acetylation and lipogenesis. Science Advances, 9(18):eadf0115, 2023.

[50] Radhakrishnan Mahadevan and Chrisophe H Schilling. The effects of alternate optimal solutions in constraint-based genome-scale metabolic models. Metabolic engineering, 5(4):264–276, 2003.

[51] Steinn Gudmundsson and Ines Thiele. Computationally efficient flux variability analysis. BMC bioinformatics, 11(1):489, 2010.

[52] Bruno G Galuzzi and Chiara Damiani. An efficient implementation of flux variability analysis for metabolic networks. In Italian Workshop on Artificial Life and Evolutionary Computation, pages 58–69. Springer, 2022.

[53] Bruno G Galuzzi, Marco Vanoni, and Chiara Damiani. Combining denoising of rna-seq data and flux balance analysis for cluster analysis of single cells. BMC bioinformatics, 23(Suppl 6):445, 2022.

[54] Sergio Bordel, Rasmus Agren, and Jens Nielsen. Sampling the solution space in genome-scale metabolic networks reveals transcriptional regulation in key enzymes. PLoS computational biology, 6(7):e1000859, 2010.

[55] Bruno G Galuzzi, Luca Milazzo, and Chiara Damiani. Adjusting for false discoveries in constraint-based differential metabolic flux analysis. Journal of Biomedical Informatics, 150:104597, 2024.

[56] Hui Zou and Trevor Hastie. Regularization and variable selection via the elastic net. Journal of the Royal Statistical Society Series B: Statistical Methodology, 67(2):301–320, 2005.

[57] James Bergstra, Daniel Yamins, and David Cox. Making a science of model search: Hyperparameter optimization in hundreds of dimensions for vision architectures. In International conference on machine learning, pages 115–123. PMLR, 2013.

[58] Leo Breiman. Random forests. Machine learning, 45(1):5–32, 2001.

[59] Tianqi Chen and Carlos Guestrin. Xgboost: A scalable tree boosting system. In Proceedings of the 22nd acm sigkdd international conference on knowledge discovery and data mining, pages 785–794, 2016.

[60] Scott M Lundberg and Su-In Lee. A unified approach to interpreting model predictions. Advances in neural information processing systems, 30, 2017.

[61] Damian Szklarczyk, Rebecca Kirsch, Mikaela Koutrouli, Katerina Nastou, Farrokh Mehryary, Radja Hachilif, Annika L Gable, Tao Fang, Nadezhda T Doncheva, Sampo Pyysalo, et al. The string database in 2023: protein–protein association networks and functional enrichment analyses for any sequenced genome of interest. Nucleic acids research, 51(D1):D638–D646, 2023.

[62] Liis Kolberg, Uku Raudvere, Ivan Kuzmin, Priit Adler, Jaak Vilo, and Hedi Peterson. g: Profiler—interoperable web service for functional enrichment analysis and gene identifier mapping (2023 update). Nucleic acids research, 51(W1):W207–W212, 2023.

[63] Fran Supek, Matko BoŠnjak, Nives Škunca, and Tomislav Šmuc. Revigo summarizes and visualizes long lists of gene ontology terms. PloS one, 6(7):e21800, 2011.

